# Preparation of PLGA Nanoparticles Encapsulated with Fluorescent Probe Coumarin-6

**DOI:** 10.1101/614875

**Authors:** Elizebeth Purr, Jacob Marshall, John Smith

## Abstract

In this report, we provided a novel platform to prepare fluorescent probe coumarin-6 nanoparticles by using biodegradable material polylactic acid-glycolic acid copolymer (PLGA) as material. The coumarin-6-PLGA nanoparticles were prepared by double emulsion and solvent evaporation. The encapsulation efficiency and releasing kinetics were also investigated. Results indicate that the encapsulation efficiency of coumarin-6 nanoparticles was 51.6%, the utilization rate was 81.9%, the average particle size was 135 nm, and the leakage rate of coumarin-6 in vitro was lower than 72 h. 2%. Our experimental results provide evidence that PLGA nanoparticles can effectively encapsulate fluorescent probe Coumarin-6 and release the probe in a controlled manner.

## Introduction

Nano-drug system based on natural or synthetic polymer materials, with small size effect, surface effect and interface effect, can cross the blood-brain barrier, reticuloendothelial tissue, and change the distribution of drugs in the body; The effect gathers and stays in the tumor site, improves the curative effect, reduces the adverse drug reaction; can improve the stability of the polypeptide, the vaccine drug in vivo, regulate the release rate, increase the permeability of the biofilm such as the gastrointestinal tract, and make it become One of the most active research directions is increasingly being used to control drug release and targeted drug delivery systems. In particular, biodegradable polymers typified by polylactic acid (PLA) and poly(lacticcoglycolic acid) (PLGA) have good biodegradability and biocompatibility. It is widely used in the field of biomedical engineering materials and microparticle drug delivery systems [1–2]. An important part of the in vivo evaluation of microparticle delivery systems is the study of their absorption and transport in vivo. The current method is to perform in vivo transport of the tracer delivery system after fluorescent labeling or radiolabeling. Since the fluorescent labeling method has no radioactive contamination problem, the in vivo transport of the drug delivery system can be visually and qualitatively observed by a fluorescence microscope or a confocal microscope, and a quantitative measurement can be performed, thereby obtaining a wider application. Fluorescent labeling methods used in laboratories include chemical synthesis [3] and physical entrapment [4]. Coumarin-6 is a kind of fat-soluble laser dye with high laser conversion rate and stable performance. The literature reports that nanoparticles containing 0.05% coumarin-6 can be observed under the microscope. Strong fluorescence, and can be quantitatively determined by HPLC, the detection sensitivity is very high [5]. Therefore, in recent years, coumarin-6 has been used as a fluorescent probe in a microparticle drug delivery system to conduct in vivo tracking, cell uptake, and transport mechanism studies of drug delivery systems [6–7]. In this experiment, the PLGA nanoparticles coated with coumarin-6 were prepared by the preparation method of PLGA nanoparticles commonly used in the laboratory-emulsification-solvent evaporation method. The orthogonal test was used to optimize the formulation process, and the morphology of the nanoparticles was investigated. The particle size distribution and the in vitro leakage of coumarin-6 in the nanoparticles, the feasibility of coumarin-6 as a fluorescent marker for PLGA nanoparticles, and the further study on the in vivo transport and targeting of drug-loaded nanoparticles basis.

## Materials and Method

RF-5301 Fluorescence Spectrophotometer (SHIMADZU, Japan); LC-10AT High Performance Liquid Chromatograph (RF-10AXL Fluorescence Detector, SHIMADZU, Japan); SZCL-2 Digital Intelligent Temperature Controlled Magnetic Stirrer (Gongyi City Yuhua Instrument) Limited company) SCIENTZ-IID ultrasonic cell pulverizer (Ningbo Xinzhi Biotechnology Co., Ltd.); JEM-100CXII transmission electron microscope (Japan Electronics Co., Ltd.); Mastersizer 2000 laser particle size analyzer (Malay, UK). Coumarin-6 (Sigma-Aldrich, USA ≥98.0%); PLGA (LA GA:=75:25, Mr 10 000, Shandong Institute of Medical Devices); Sephadex G-50 Sephadex (Sweden) Pharmacia); polyvinyl alcohol (PVA 4-88, Mr 31 000, Sigma-Aldrich, Germany); methanol (chromatographically pure, Merck, Germany); methanol, acetonitrile, dichloromethane, ethyl acetate (analytical grade, Tianjin Fuyu) Fine Chemical Co., Ltd.)

## Restuls

### Preparation of coumarin-6 PLGA nanoparticles

The PLGA nanoparticles were prepared by emulsification-solvent evaporation method. The appropriate amount of PLGA and coumarin-6 were dissolved in a certain amount of dichloromethane and ethyl acetate (7:3) as organic. Phase, add appropriate amount of PVA aqueous solution, in an ice water bath condition, use ultrasonic cell pulverizer intermittent ultrasonic for several minutes, transfer to 0.5% PVA aqueous solution, stir on a magnetic stirrer for 3 ~ 4 h, remove the organic solvent, get Coumarin-6-PLGA nanoparticles. Blank nanoparticles containing no coumarin-6 were prepared by the same method.

### HPLC analysis

Specificity test precision weighed 5.09 mg of coumarin-6 in a 50 mL volumetric flask, dissolved in methanol and brought to volume as a coumarin-6 stock solution. Accurately absorb the appropriate amount of coumarin-6 stock solution, add methanol to volume as a reference solution; accurately absorb the appropriate amount of nanoparticles containing coumarin-6, add acetonitrile to dissolve the material, and finally add appropriate amount of methanol to dilute, as the test solution; Another appropriate amount of blank nanoparticles was taken, and a negative control solution was prepared by the same method. A 20 μL assay was injected in sequence and the chromatogram was recorded. The results showed that the retention time of coumarin-6 was about 5.6 min and the peak shape was good. The separation of coumarin-6 from the adjacent impurity peaks in the test sample was greater than 1.5, and the theoretical plate number was calculated according to the peak of the control product >3 000. The negative control solution did not interfere with the determination, indicating that the analytical method has good specificity. The chromatogram is shown in Figure 1.

**Figure 1.**
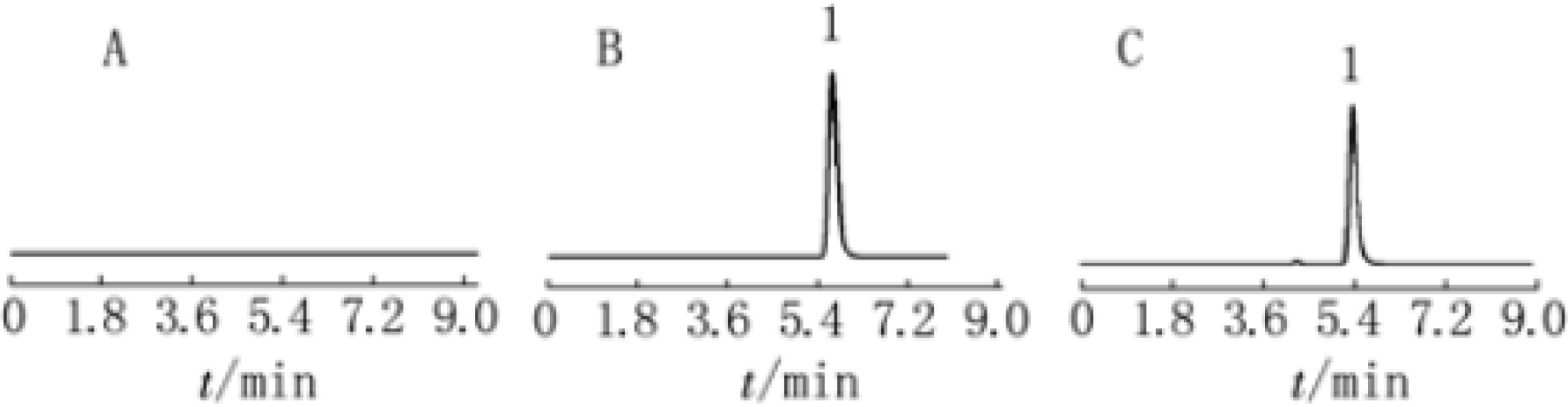
HPLC analysis

### Recovery test

Take 0.1 mL of blank nano-colloidal solution, and add the appropriate amount of coumarin-6 stock solution, dissolve the material with appropriate amount of acetonitrile, and dilute with methanol to prepare three kinds of solutions of low, medium and high concentrations. Inject 20 μL of the assay, record the peak area, substitute the linear equation, and calculate the recovery. Results The recoveries of different concentrations of coumarin-6 ranged from 98% to 104%, and the RSD value was 1.88%, indicating that the recovery rate of this method is good, which meets the requirements for in vitro sample determination.

### Encapsulation Efficiency

The coumarin-6-loaded nanoparticles and free coumarin-6 were separated using a Sephadex column. Sephadex G-50 packed in a column (l.5 cm × 10 cm) was swelled for more than 24 h, and 1 mL of the colloidal liquid was accurately sampled and eluted with deionized water to collect the separated coumarin-6. Nanoparticles. Accurately absorb 0.1 mL of coumarin-6 nanoparticle and 0.5 mL of coumarin-6 nanoparticle separated by column, add appropriate amount of acetonitrile to dissolve the material, and then dilute with appropriate amount of methanol Filtration of membranes, injection of 20 μL, to obtain the total amount of coumarin-6 in the nanoparticle system WT and encapsulated in nanoparticles The amount of coumarin-6 in the WNP. In addition, W0 is the total input of coumarin-6, and the encapsulation efficiency and utilization rate of coumarin-6 are calculated according to the following formula:

### Orthogonal Optimization

On the basis of the pre-test, the concentration of PLGA, the volume ratio of internal and external phase, the concentration of PVA and the ratio of drug to carrier material were selected as the investigation factors. The experiment was arranged according to the orthogonal design L9(34) table to encapsulate the rate and utilization rate. In order to optimize the indicators, the main influencing factors and the optimized prescriptions were determined according to the analysis of variance. The table of each factor level is shown in Table 2. The test results are shown in Table 3, Table 4 and Table 5. The results of orthogonal analysis showed that the encapsulation efficiency of coumarin was optimized. The order of importance of various factors was the ratio of internal to external phase volume > PLGA concentration > PVA concentration > ratio of coumarin to PLGA; Taking the utilization ratio of coumarin as the optimization index, the ratio of coumarin to PLGA is > PLGA concentration > PVA concentration > internal and external phase volume ratio. It indicates that the volume ratio of inner and outer phase has a great influence on the encapsulation efficiency, while the ratio of feed to material has a great influence on the utilization rate of coumarin-6. The encapsulation efficiency is an important indicator for the quality evaluation of nanoparticles. The high encapsulation ratio indicates that the proportion of coumarin-6 encapsulated in the nanoparticles is high; and the utilization rate is the total amount and input of coumarin-6 in the nanoparticle system. The higher the utilization rate, the higher the amount of coumarin-6 entering the nanoparticle system, and the smaller the loss during the preparation process^1^, which is also a non-negligible indicator for evaluating the nanoparticle formulation process. Therefore, the two factors are considered comprehensively, and the results are analyzed with comprehensive indicators (encapsulation ratio × 0.7 + utilization × 0.3). The results of variance analysis showed that the ratio of coumarin to PLGA, the ratio of internal and external phase volume and PLGA concentration had significant effects on the preparation of nanoparticles. The final optimized formulation is D3B2A3C3, which is PLGA 30 mg⋅m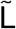1, the oil-water volume ratio is 1:5, the PVA concentration is 3%, the coumarin to PLGA weight ratio is 1:500, 51.6%, and the drug loading is 0.08%.

### Morphology Characterization

Take coumarin-6 PLGA nanoparticles in appropriate amount with 1.5% phosphotungstic acid Negative dyeing, dripping on the coated electron microscope copper net, dried and placed^2–12^. The morphology of the nanoparticles were observed under transmission electron microscope. Use The particle size distribution was measured by a laser particle size analyzer. The results are shown in Figure 2 and 3. The photo shows that the nanoparticles are round and the size is relatively uniform. There is basically no adhesion between the rice grains. The particle size distribution is narrow and concentrated It is distributed in the range of 100~190 nm with an average particle size of 135 nm.

**Figure 2.**
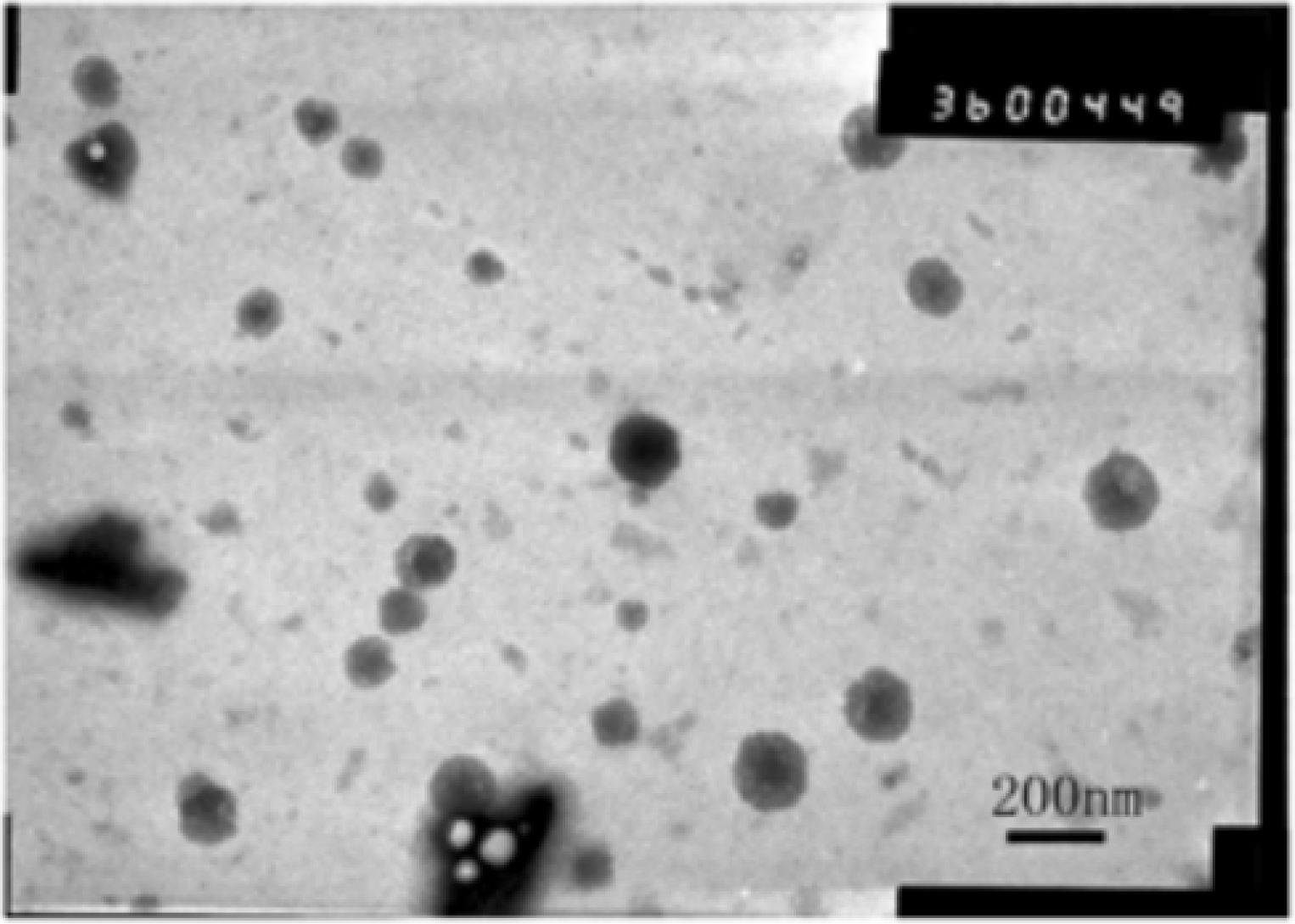
Assess PLGA nanoparticle morphology by utilizing transmission electron microscopy.

**Figure 3.**
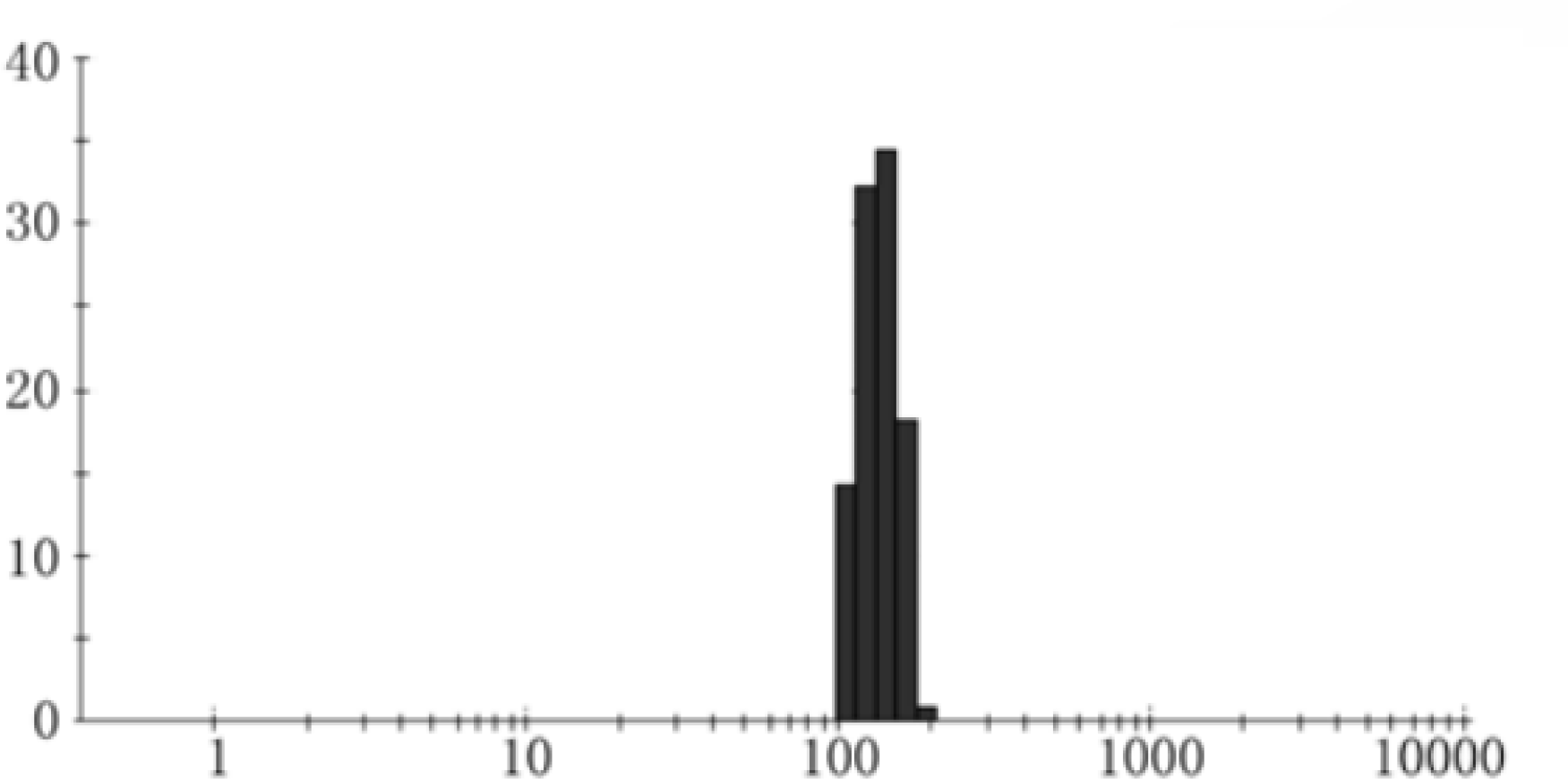
PLGA nanoparticle size and dispersity was analyzed by dynamic light scattering.

### Releasing Kinetics

Precisely measured the coumarin −6 nanoparticle glue after the column 3 mL, placed in a pre-treated dialysis bag, adding pH 7.4 of 0.1 mol.L in 2.0 mL of phosphate buffered saline (PBS). Both ends of the bag were placed with a plug cone containing 15 mL of phosphate buffer In the bottle, placed in a constant temperature oscillator, temperature 37 °C, oscillation frequency 100 r/min at 0.5, 1, 2, 4, 8, 24, 48, respectively. Take 1 mL of release medium at 72 h and immediately replenish the release medium at the same temperature. 1 mL. The concentration of coumarin-6 was determined by HPLC, and the cumulative release was calculated. The fraction is plotted and the release curve is plotted. The results are shown in Figure 4. Result coumarin-6 The cumulative release percentage at 72 h is <2%, indicating that the fluorescent substance is capable of The stable package is contained in the nanoparticles, which is not easy to leak and has no sudden release. It can be used as a fluorescent marker to trace the in vivo rotation of nanoparticles more accurately.

**Figure 4.**
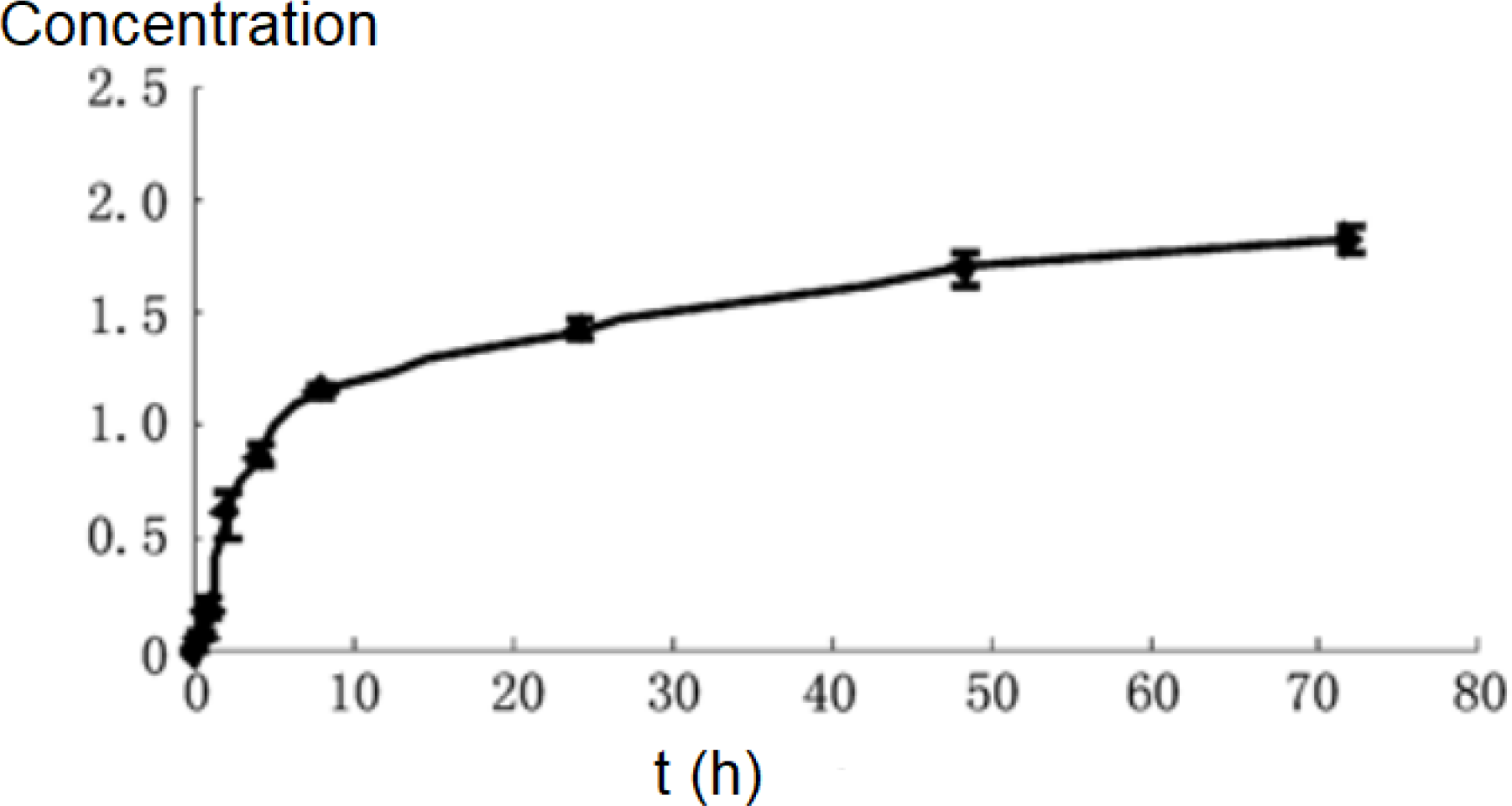
Releasing kinetics of fluorescence probe from PLGA nanoparticles.

## Discussion

PLGA nanoparticles were prepared by emulsification-solvent evaporation method. The physical properties of organic solvents (such as surface tension, vapor pressure, etc.) have a significant effect on the formation of nanoparticles [10]. The effect of the ratio of dichloromethane and ethyl acetate in the organic phase on the preparation of nanoparticles was investigated. The results were obtained by preparing coumarin-6 nanoparticles with an organic solvent of dichloromethane:ethyl acetate of 7:3. A product with a smaller particle size. Orthogonal tests showed that the ratio of coumarin to carrier materials, the ratio of internal and external phase volume, and PLGA concentration had a significant effect on the preparation of coumarin-6 nanoparticles. PVA has an emulsification and stabilization effect. A concentration of more than 1% can prepare a relatively stable emulsion, and its concentration has little effect on the encapsulation efficiency of coumarin-6. For fluorescent labeling by physical entrapment, the basic requirement for the in vivo transport behavior of the tracer nanoparticles is that the label entrapped in the nanoparticles during in vivo transport remains stable and does not leak. The in vitro leak test is an important method to investigate the feasibility of coumarin-6 as a fluorescent probe^1, 13–14^. The percentage of leakage of coumarin-6 nanoparticles in buffer for 72 h was measured.

## Conclusion

The preparation process of coumarin-6 PLGA nanoparticles is simple and feasible. Coumarin-6 can be used as a fluorescent probe for the tracer and targeting of nanoparticles in animals. Future efforts will focus on translating this promising technology in a clinical setting.

